# Single-cell resolution map of innate-like lymphocyte response to *Francisella tularensis* infection reveals MAIT cells’ role in protection from tularemia-like disease

**DOI:** 10.1101/2024.10.04.616759

**Authors:** G. Donald Okoye, Amrendra Kumar, Farshad Ghanbari, Nowrin U. Chowdhury, Lan Wu, Dawn C. Newcomb, Luc Van Kaer, Holly M. Scott Algood, Sebastian Joyce

## Abstract

Early immune dynamics during the initiation of fatal tularemia caused by *Francisella tularensis* infection remain unknown. Unto that end, we generated a transcriptomic map at single-cell resolution of the innate-like lymphocyte responses to *F. tularensis* live vaccine strain (LVS) infection of mice. We found that both interferon-γ-producing type 1 and interleukin-17-producing type 3 innate-like lymphocytes expanded in the infected lungs. Natural killer (NK) and NKT cells drove the type 1 response, whereas mucosal-associated invariant T (MAIT) and γδ T cells drove the type 3 response. Furthermore, tularemia-like disease-resistant NKT cell-deficient, *Cd1d*^-*/-*^ mice accumulated more MAIT1 cells, MAIT17 cells, and cells with a hybrid phenotype between MAIT1 and MAIT17 cells than wild-type mice. Critically, adoptive transfer of LVS-activated MAIT cells from *Cd1d*^-*/-*^ mice, which were enriched in MAIT17 cells, was sufficient to protect LVS-susceptible, immunodeficient *RAG2^-/-^* mice from severe LVS infection-inflicted pathology. Collectively, our findings position MAIT cells as potential mediators of interleukin-17-dependent protection from pulmonary tularemia-like disease.

**HIGHLIGHTS:**

- Pulmonary *F. tularensis* LVS infection induces type 1 & type 3 immune responses.
- NK and NKT cells drive type 1, whilst MAIT and γδT cells drive type 3 responses.
- Increased MAIT17 cell accumulation is associated with resistance to TID.
- Adoptive transfer of MAIT17-enriched cells protect immunodeficient mice from TID.

## INTRODUCTION

Barrier mucosae form vulnerable habitats for infections. Resident myeloid cells and subsets of unconventional lymphocytes—which include natural killer T (NKT), mucosal-associated invariant T (MAIT) and γδ T cells—collectively called innate-like T lymphocytes (ITLs), continuously surveil the barrier mucosae, including those of the lung, forming the first line of defense against pathogenic agents. The Gram-negative intracellular bacterium *Francisella tularensis*, the etiologic agency of tularemia, is one such pathogen. Natural *F. tularensis* infections are currently endemic in many parts of Europe with sporadic outbreaks reported in the United States^1–3^. Natural *F. tularensis* infection acquired by inhalation frequently causes fatal pulmonary tularemia^4, 5^. Infection following ingestion of *F. tularensis* causes gastrointestinal disease, whereas transmission by blood-sucking insects, such as fleas, ticks, and deer flies, as well as exposure of open wounds to the bacterium cause ulceroglandular tularemia^4, 6, 7^. Natural infections when diagnosed early are curable by antibiotics^7^. When untreated, *F. tularensis* organisms from ulceroglandular lesions can disseminate, induce sepsis, and result in pulmonary tularemia^4, 5, 7, 8^. Moreover, respiratory infections with as few as ten aerosol droplets containing type A *F. tularensis* subspecies tularensis cause fatal pulmonary tularemia^6, 8^. This high infectivity has raised type A *F. tularensis* to the top of tier 1 select agent list of pathogens of high interest^9^. The live vaccine strain (LVS) derived from type B *F. tularensis* subspecies holarctica can protect against the virulent type A *F. tularensis*, but only when administered via the respiratory route^10–12^, implicating lung-resident immune cells in protective immunity. LVS is not yet licensed as a tularemia vaccine by the FDA, as its administration via the respiratory route is associated with adverse effects^13^. Hence, there is an unmet need to understand the immune mechanisms of protection to inform targets for the design of better vaccines and therapeutics against tularemia.

Early immune dynamics during the initiation of fatal tularemia remain largely unknown. Current evidence suggests that murine and human natural killer (NK) cells, which primarily produce interferon-γ (IFN-γ) in response to LVS infection in a MyD88-dependent manner, are critical for bacterial clearance^14–16^. Activation of mouse hepatic NK cells during LVS infection reduces hepatic cell death, induces granuloma formation, and limits hepatic necrosis^17^. However, the NK cell response during LVS infection is redundant, as other immune cells, such as myeloid lineage and adaptive immune cells, can compensate for the absence of NK cells^18, 19^.

NKT cells are critical early responders to bacterial invasion^20^. NKT cells patrol common sites of microbial entry including the respiratory mucosa, where they are activated by self- or non-self-glycolipids presented by CD1d molecules^21, 22^. Indeed, B cell-NK cell-NKT cell crosstalk in mice can control *F. tularensis* infection^23^. By contrast, we previously showed that the NKT cell response is detrimental for survival from LVS infection, as NKT cell-deficient *Cd1d^-/-^* mice are resistant to tularemia-like disease (TLD)^24^. The *Cd1d^-/-^* mice exhibit far less type 1-driven sepsis-like inflammation, consistent with runaway inflammation being a hallmark of tularemia^24, 25^. Similarly, in other models of intracellular bacterial infections, NKT cells failed to protect mice from *Mycobacterium tuberculosis* infection and were detrimental in *Legionella pneumophilia* and *Chlamydia trachomatis* infections where they drive a type 1 inflammatory response^26–28^. As NKT cells have previously been implicated in sepsis, their presence may exacerbate inflammation and, thus, impair survival of infected mice^29, 30^. Notably, LVS-infected *Cd1d^-/-^* mice showed increased levels of induced bronchus-associated lymphoid tissues (iBALTs), which have been considered the basis of protection in this mouse model^24^. How a single gain-of-function mutation—i.e., CD1d-deficiency, confers resistance to TLD remains unclear.

MAIT cells, akin to NKT cells are also found at sites of microbial entry. They are essential for protective responses to LVS infection in mice similar to other intracellular bacterial infections including *L*. *longbeachae*, *M*. *tuberculosis*, and *M*. *abscessus*^31–35^. MAIT cells are activated by the presentation of vitamin B2 metabolites on MHC-related MR1 molecules^36–38^. MAIT cell activation is primarily dependent on TCR-mediated activation, but cytokine receptors may also contribute to MAIT cell activation and function during infection^32, 39–41^. Importantly, there are two distinct subsets of MAIT cells in mice and humans: the type 1 MAIT1 and type 3 MAIT17 which cytokines that mirror T helper 1 (Th1) and Th17 responses, respectively^42, 43, 44^. They are metabolically distinct and play protective roles in infectious diseases^43^. Recent evidence suggests that MAIT1 cells are essential and sufficient to protect against systemic LVS infection^44^. What roles MAIT17 cells play in pulmonary LVS infection, which causes lethal tularemia, is unknown.

Human γδ T cells control both LVS and virulent type A *F. tularensis* SchuS4 growth in vitro in an IFN-γ-dependent manner^45^. By contrast, protection against pulmonary LVS infection in both humans and mice requires interleukin-17 (IL-17)^46–50^. As γδ T cell-deficiency caused lethal TLD in mice, this T cell subset was considered the relevant source of IL-17^48, 51^. By contrast, γδ T cells play a detrimental role in cutaneous *F. tularensis* LVS infection^51^. However, intranasal infection of mice with LVS induces a TCR-dependent expansion of IL-17-producing γδ T cells, important for adequate bacterial clearance^48, 52^. Viewed together, current evidence suggests that NK, NKT, MAIT and γδ T cells as well as IFN-γ and/or IL-17 secreted by them are protective or detrimental to LVS infections. Nonetheless, the dynamics of protective innate-like lymphocyte responses to LVS infection remain unclear.

To gain insights into the questions raised above, we focused on a mouse model of pulmonary tularemia because it is the most lethal form of *F. tularensis* infection. In this preclinical model, intranasal inoculation with LVS induces pulmonary TLD. We reasoned that correlates of immune protection will emerge from a comparative study of TLD-sensitive C57BL/6 and TLD-resistant *Cd1d^-/-^* mice. Hence, a transcriptome profile was established at single-cell resolution to understand the dynamics of lymphocyte responses in the lungs of C57BL/6 and *Cd1d^-/-^* mice during the acute phase of LVS infection. Then, the key findings were validated by complementary immunoassays. Finally, the ability of MAIT cells alone to transfer protection in immunodeficient mice was interrogated. Collectively, our findings indicated that, in addition to an anticipated IFN-γ-producing type 1 response, which occurs first, respiratory LVS infection also induced a robust IL-17-producing type 3 response driven in large part by lung-resident MAIT17 cells. These lung-resident MAIT cells are sufficient to confer protection to LVS-susceptible, immunodeficient mice from fatal TLD.

## RESULTS

### Respiratory LVS infection elicits type 1 & type 3 ITL responses

To understand the dynamics of the immune response to respiratory LVS infection during the critical stage of tularemia-like disease (TLD), six-week-old male C57BL/6 mice were intranasally inoculated with PBS (mock) or ∼9,000 cfu LVS—the LD_50_ previously established^24^. Seven days post inoculation (dpi), CD45^+^ hematopoietic cells were flow sorted from single cell suspensions prepared from whole lung tissue of mock and LVS-infected mice. The B cell-negative cell fraction (B220^+^ cells electronically gating out) of >98% viability (**Figure S1A**) was subjected to scRNAseq using 10x Genomics platform (see Methods). Quality control (as described in the Methods) yielded a total of 13,063 high quality and comparable cells across the mock and LVS-infected samples. While we attempted to deplete CD14^+^ cells by flow sorting, CD14 staining was ineffective as CD11b^+^ & CD11c^+^ myeloid cells remained detectable in substantial numbers (**Figure 1A**). Because the incomplete depletion of CD14^+^ cells would undermine any analyses of the myeloid compartment, we focused on the B cell negative, CD45^+^ lymphocyte compartment. Within this population, both adaptive (tagged naïve and effector T) and innate/innate-like (tagged innate-like T and NK cells) lymphocytes were evident. Each cluster was identified and annotated using the scType computational platform for scRNAseq data^53^ and confirmed by manual verification of signature genes as described in the Methods section. Surprisingly, while a conventional T cell response to LVS infection at 7 dpi was evident, CD4^+^ and CD8^+^ T cell expansion was not as robust (**Figure 1A**) as would be expected from the critical role they are thought to play to protect against LVS infection^54–56^. Just as surprisingly, only NK cells in the innate lymphoid cell lineage were detectable in this experiment (**Figure 1A**). Hence, in the studies described below, NK cells were analyzed together with unconventional T cells.

**Figure 1:**
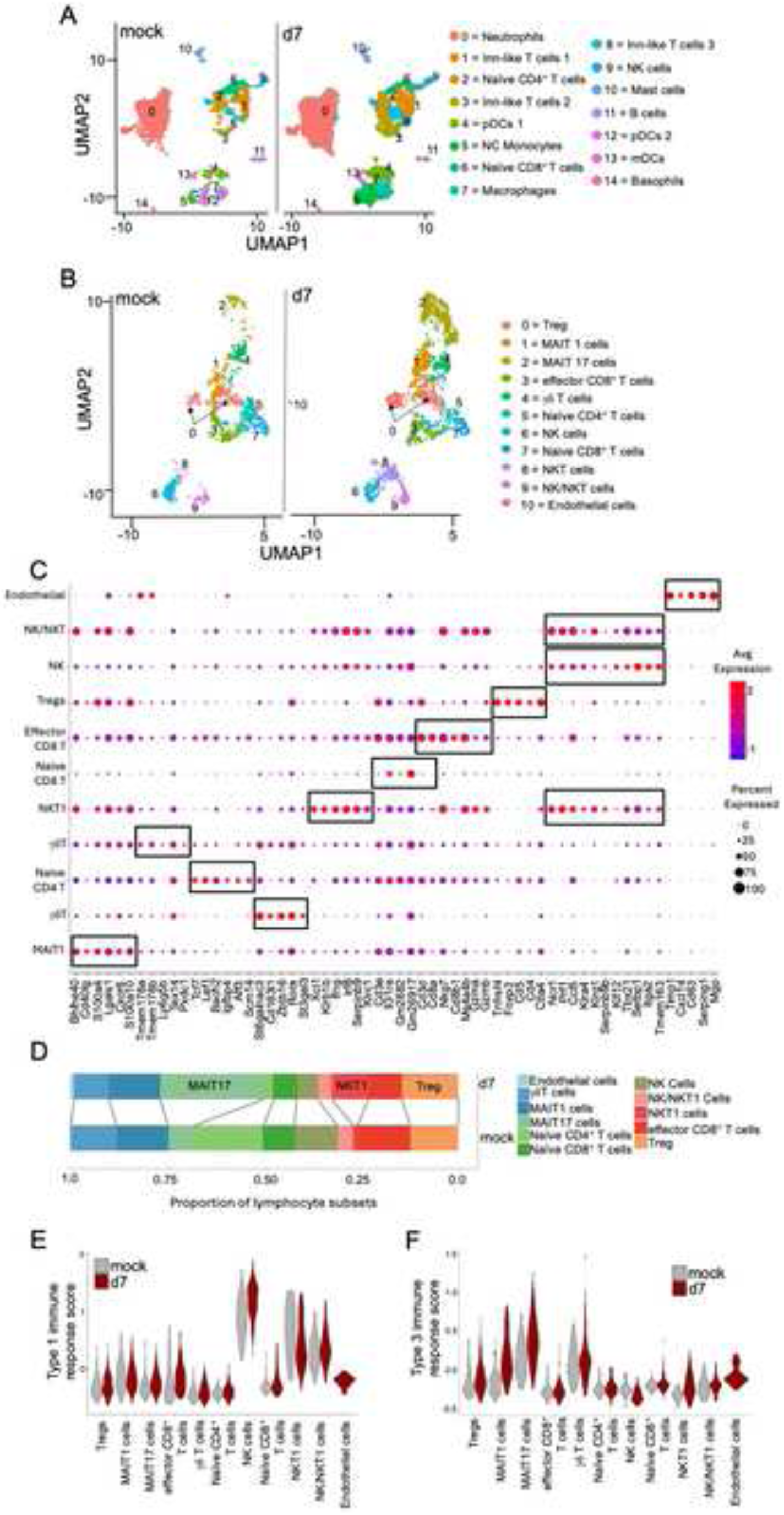
Intranasal LVS infection of C57BL/6 mice expands myeloid cells and innate-like lymphocytes in the lungs. Three-four six-week-old C57BL/6 mice were inoculated i.n. with PBS (mock) or ∼9,000 cfu (previously calculated lethal dose (LD_50_ = 8,100 cfu) that kills ∼50% of C57BL/6 mice) LVS. At 7 dpi (days post inoculation), the B220^+^ B cell and CD14^+^ myeloid cell negative fraction of CD45^+^ cells were flow sorted from single cell suspensions prepared from whole lung tissue of mock or LVS-infected mice. Flow sorted cells >98% viable (see Figure S1A for gating strategy) were subjected to scRNAseq (see Methods). **A.** Annotated uniform manifold approximation and projection (UMAP) visualization of 13,063 hematopoietic cells from the lungs of mock and LVS-infected mice. Fifteen clusters were identified featuring cells of the innate, innate-like and adaptive immune system. Note that staining with anti-CD14 mAb did not deplete myeloid cells in this experiment. **B.** UMAP plot of T cells clustered by identity and separated by condition. For this, T cells (2,598) from **A** were extracted and re-clustered for deeper analysis of innate and innate-like lymphocytes. **C.** Dot plot showing the scaled mean expression of marker genes that were used to identify cell types to cell clusters. The color intensity represents average expression of each marker gene in each cluster. The dot size indicates the proportion of cells expressing each marker gene. The full gene list of markers for cluster identification is provided in Supplementary Table 1. **D.** Proportion of lymphocyte subsets by cluster in mock infected and at 7 dpi. **E.** Violin plot for scaled expression of type 1 immune response gene signature [*Tbx21, Ifng, Gzma, Gzmb, Tnf, Bhlhe40, Klrb1c, Klra8, Prf1, Cx3cr1, Il2rb, Klre1,* and *Ccr5* ^57^]. **F.** Violin plot for scaled expression of type 3 immune response gene signature [*Il17a, Il17f, Ccr6, Rorc, Irf4, Il23r, Il1r1, Lmna, Tnf, Nebl,* and *Lamc1* ^58^. Data for **A—E** were generated from pooled samples of 3 mice for infected and 4 mock-treated mice.

In silico sorting of cells from each sample allowed further analyses of lineages of interest. Hence, to focus on and characterize NK and unconventional T cell responses to acute LVS infection of the lungs in detail, 2,598 cells assigned to the lymphocyte clusters in the above experiment were extracted by in silico sorting, re-clustered and analyzed. Each cluster was identified by gene signatures (**Table S1 & Figure 1B&C**) as described in the Methods section. The most obvious expansion identified unconventional, innate-like T lymphocytes (ITLs), which included NKT and MAIT cells (**Figure 1B—D**). Further, Treg cells also expanded and a concomitant reduction in other lymphocytes including NK cells was observed (**Figure 1B—D**).

We next defined ITL subsets activated during respiratory LVS infection. This analysis identified ITL clusters expressing *Ifng* and *Il17a*, and their associated transcription factors, implying the presence of at least two inflammatory response profiles (**Figure S1B&C)**. Thus, to establish the specific types of inflammatory responses associated with LVS infection, all lymphocyte clusters were examined for expression of genes associated with type 1 and type 3 inflammation adapted from previously published gene sets^57, 58^. Type 1 immune signature genes included *Ifng, Tbx21, Gzma, Gzmb, Tnf, Bhlhe40, Klrb1c, Klra8, Prf1, Cx3cr1, Il2rb, Klre1,* and *Ccr5* ^57^, whereas type 3 genes included *Il17a, Il17f, Rorc, Ccr6, Irf4, Il23r, Il1r1, Lmna, Tnf, Nebl,* and *Lamc1* ^58^. This analysis revealed that the response was characterized by an NK and NKT cell-driven type 1 and a MAIT and γδ T cell-driven type 3 response (**Figures 1E&F and S1D&E**). Thus, pulmonary LVS infection triggers both type 1 and type 3 inflammatory responses.

### Respiratory LVS infection of mice elicits ITL responses 5 days post inoculation

The immune response to respiratory LVS infection begins approximately 72 hours post inoculation^14, 24, 25, 44, 59^. This route of infection induces prostaglandin E2 production in the lungs which in turn promotes the release of anti-inflammatory factors. These anti-inflammatory factors inhibit the generation of IFN-γ-producing CD4^+^ T cells ^60, 61^. Whether this capacity for host immune evasion by LVS infection extends to ITL responses is unknown. Hence, the dynamics of the ITL response was evaluated in a time course study. In this experiment, C57BL/6 mice were inoculated with PBS (mock) or ∼8,500 cfu LVS by the intranasal route as above, and the response evaluated at 3, 5 and 7 dpi. At these time points, to improve lymphocyte yields, CD11b^+^ and CD11c^+^ myeloid as well as B220^+^ B cells were removed by flow sorting and the remaining CD45^+^ cells were subjected to scRNAseq as above. Quality control yielded a total of 8,326 high quality and comparable cells across all four samples (**Figure 2A**). Consistent with findings in conventional T cells, we observed NK and ITL accumulation after infection with significant expansion only from 5 dpi onward (**Figure 2B&C**). Moreover, type 1 and type 3 inflammatory responses were driven by NK, NKT1 and MAIT1 cells and by MAIT17 and γδ T cells, respectively (**Figure 2D&E and S2A-D**). Furthermore, type 1 immune response peaked by 5 dpi while type 3 immune response continued to increase until at least 7 dpi (**Figure 2D&E**), suggesting a differential role for these different ITL subsets in the immune response to respiratory LVS infection. Taken together, these data support a model in which respiratory LVS infection of mice expands both type 1 and type 3 ITLs and attendant inflammatory responses only by 5 days post inoculation.

**Figure 2:**
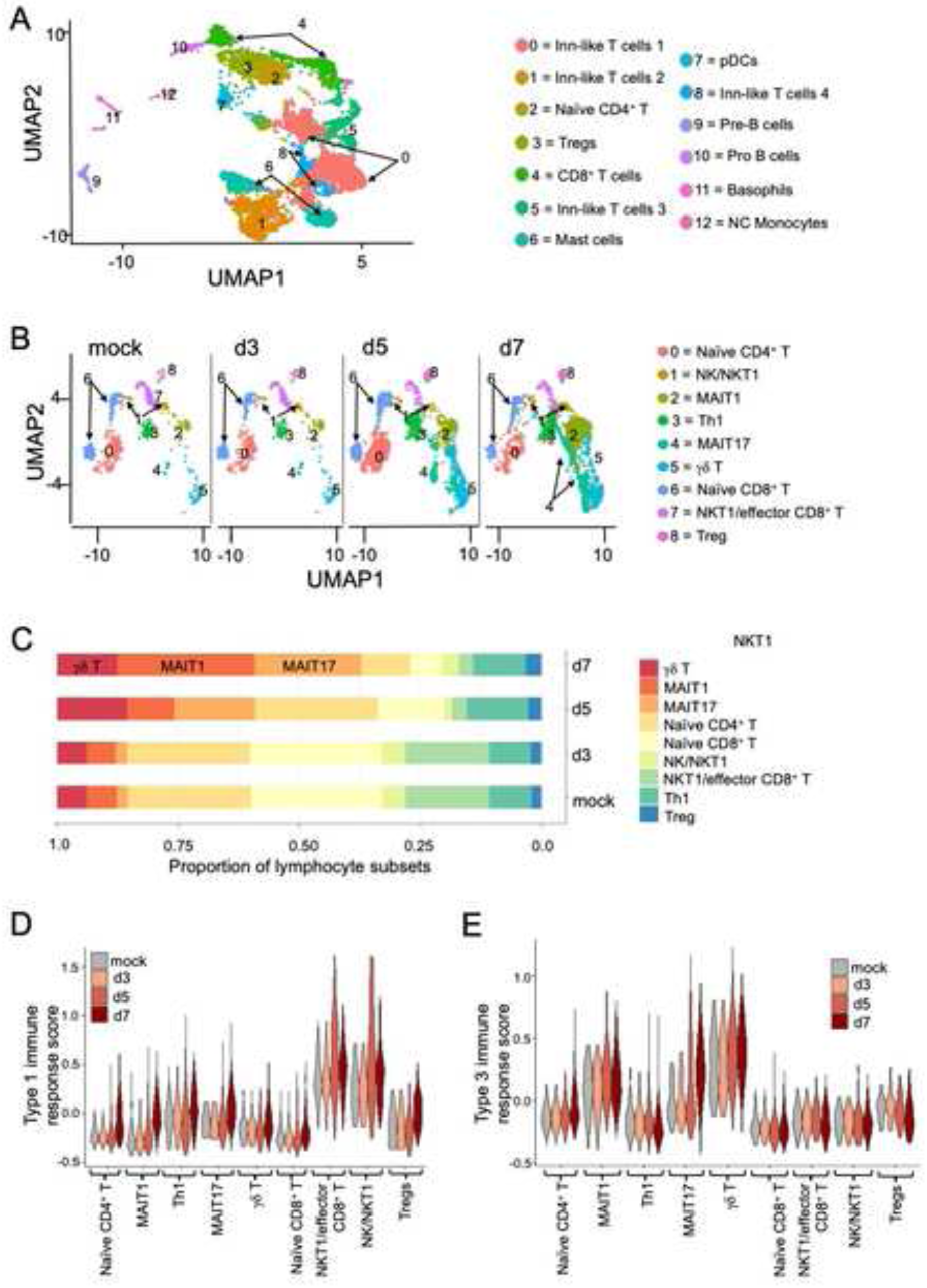
LVS infection of C57BL/6 mice elicits type 1 and type 3 inflammatory responses from innate-like T lymphocytes. Three six-week-old C57BL/6 mice were inoculated by the i.n. route with PBS (mock) or ∼8,500 cfu LVS. At 3, 5, and 7 dpi, the B220^+^ B cell as well as CD11b^+^ and CD11c^+^ myeloid cell negative fraction of CD45^+^ cells were flow sorted from single cell suspensions prepared from whole lung tissue of mock or LVS-infected mice. Flow sorted cells with >98% viability were subjected to scRNAseq (see Methods). **A.** Annotated UMAP plot of 8,326 lung CD45^+^ cells from mock and d3, d5 and d7 LVS-infected mice are shown. B220^+^ B cells as well as CD11b^+^ and CD11c^+^ myeloid cells were excluded from CD45^+^ cells during flow sorting. Clusters were inferred by cluster identity and color-coded as shown. **B.** UMAP plots of lung T cells colored by inferred cluster identity are represented for mock and d3, d5 and d7 LVS-infected mice. For this, 6,724 lymphocytes were extracted in silico from A and re-clustered for further analyses. **C.** Proportion of lymphocyte subsets by cluster per condition (mock, 3, 5 and 7 dpi). **D.** Violin plot for scaled expression of type 1 immune response gene signature based on genes listed in figure 1E legend. **E.** Violin plot for scaled expression of type 3 immune response gene signature based on genes listed in figure 1F legend. Data for A—E were generated from pooled samples of 3 mice per condition.

### NK and NKT1 cells contribute to type 1 immune responses in LVS-infected C57BL/6 lungs

We previously reported that NKT cell-deficient *Cd1d^-/-^* mice are relatively resistant to respiratory LVS infection-induced TLD^24^. This was associated with a reduced inflammatory milieu in *Cd1d^-/-^* mice when compared to C57BL/6 mice. Hence, an outsized type 1 response driven by NKT cells may underlie the reported lung pathology to LVS infection in C57BL/6 mice^24^. Notably, NK cells contribute significantly to type 1 inflammation in response to LVS infection^62, 63^. Thus, to explore NK and NKT cell involvement, all NK, NKT and associated cell clusters were sorted in silico and sub-clustered for additional analysis (**Figure 3A**). Sustained NK cell presence was observed with notable expression of type 1 inflammatory genes including *Ifng* (**Figure 3B—D**). An early expansion of NKT1 cells by 5 dpi expressing *Ifng* and *Tbx21*, which encodes Tbet—a Th1-typic transcription factor (TF) that drives type 1 responses, was also observed (**Figure 3B—D, S3A&B**). *Ifng* transcript was apparent in NK and NKT1 cells from uninfected mice, as was previously reported^64^. The *Ifng* transcripts peaked by 5 dpi and persisted until at least 7 dpi (**Figure 3D**). Importantly, neither NKT2 nor IL-17-producing NKT17 cell responses were observed (**Figures 3E&F, S3C-E**). Accordingly, flow cytometric evidence indicated that LVS infection induces a rapid NKT1 cell expansion with a concomitant NKT2 and NKT17 subset contraction in both the lungs and spleen (**Figure S3F**).

**Figure 3:**
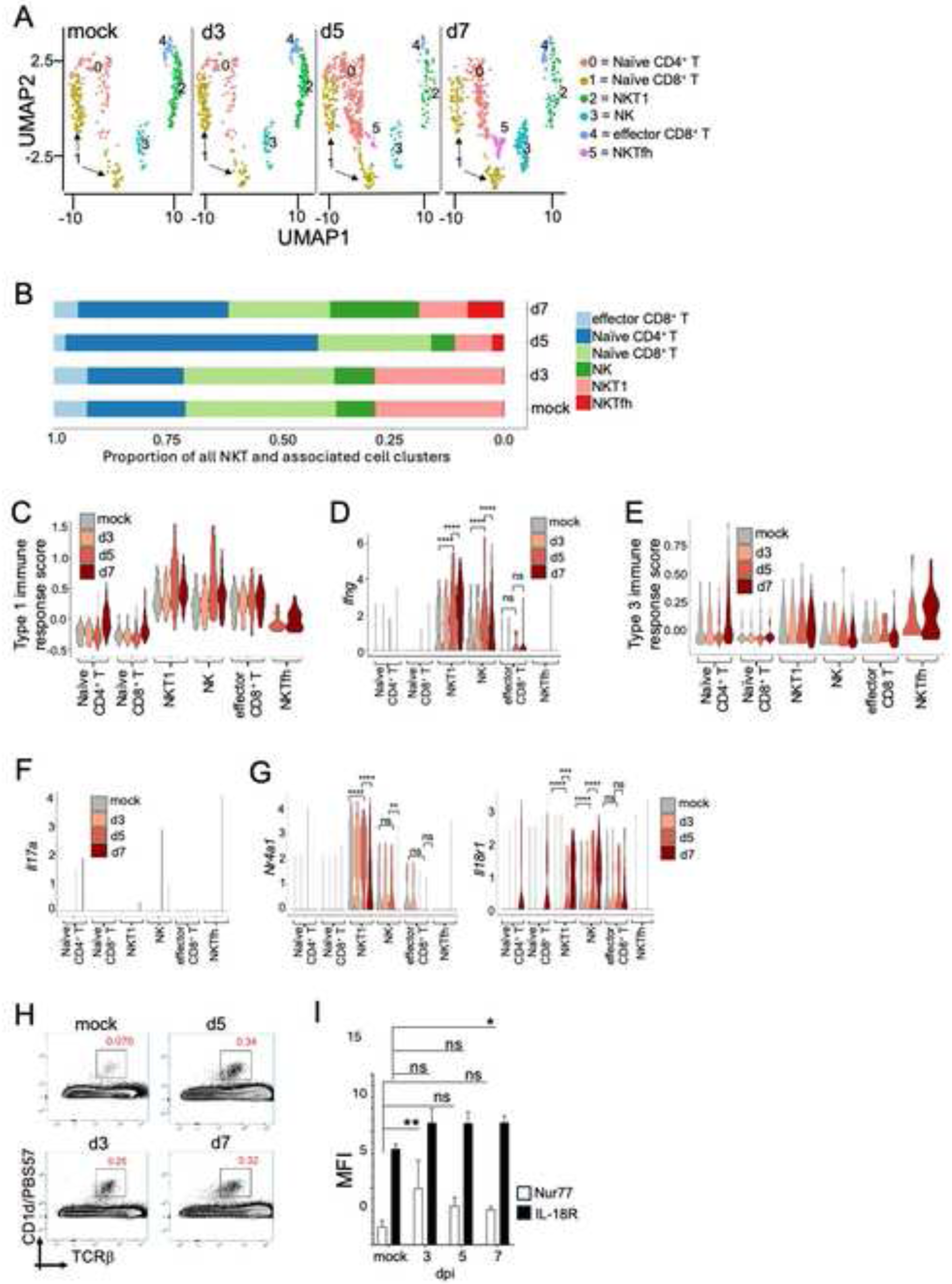
Lung NKT1 cells contribute to type 1 immune response against LVS infection in C57BL/6 mice. **A.** UMAP plot of lung type 1 cells—NKT, NK and CD8^+^ T cells, as well as associated cell clusters color coded by cluster identity, corresponding to mock and LVS infection at 3, 5 and 7 dpi. Annotation of 1,870 type 1 naïve and effector clusters in silico extracted from figure 2B and re-clustered for further analysis. **B.** Proportion of NKT and associated cell subsets by cluster per condition. **C.** Violin plot for scaled expression of type 1 immune response gene signature based on genes listed in figure 1E legend. **D.** Violin plot for the expression of the hallmark type 1 cytokine *Ifng* in response to mock or LVS infection. **E.** Violin plot for scaled expression of type 3 immune response gene signature based on genes listed in figure 1F legend. **F.** Violin plot for the expression of *Il17a* in response to mock or LVS infection. **G.** Violin plots for *Nr4a1*, which encodes Nur77—a marker of TCR-mediated activation of NKT1 cells, and *Il18r1*—a marker of cytokine-dependent activation of NKT1 cells. Data for A—G were generated from pooled samples of 3 mice per condition. Mann Whitney test with Bonferroni posttest correction. *p<0.05, **p<0.01, ***p<0.001, ****p<0.0001. **H—I** Representative flow plots (**H**) and normalized mean fluorescence intensity (MFI; see Methods) (**I**) of Nur77 and IL18Ra expression by pulmonary NKT cells in mock or LVS infected mice. Bars represent the mean+SEM (*n*=3—5 mice per group per experiment). Two-way ANOVA with Tukey’s multiple comparison test. *p<0.05, **p<0.01, ***p<0.001, ****p<0.0001.

NKT cells can become activated by TCR engagement with glycolipid agonists, self-lipid agonists in combination with inflammatory cytokines, or inflammatory cytokines. Hence, to establish the mode of NKT cell activation by respiratory LVS infection, the expression of *Nr4a1*— which codes for Nur77 and serves as a proxy for TCR engagement, and *Il18r1*—which provides a readout for cytokine-mediated activation^65–68^, was assessed in the in silico-sorted NKT cells. An early expression of *Nr4a1* by NKT1 cells was observed, indicating initial TCR engagement (**Figure 3G**). Subsequent decrease in *Nr4a1* expression was associated with a commensurate increase in *Il18r1* expression by 7 dpi, implying continued NKT cell engagement that is cytokine-dependent but TCR-independent (**Figure 3G**). Consequently, flow cytometry experiments using Nur77 reporter mice confirmed an early and sustained accumulation of NKT cells, and revealed a significant increase in Nur77 levels in NKT cells by 3 dpi with a subsequent drop in Nur77 levels accompanied by increased IL-18Ra levels (**Figure 3H&I, S3G**). Taken together, the data indicate LVS infection induces NK cell and TCR-mediated NKT1 cell expansion that drives a type 1 inflammatory immune response.

NKTfh cells—a T-follicular-helper cell-like NKT cell subset, also were identified. These cells alongside effector CD8^+^ T cells also appear to elicit a type 1 immune response to respiratory LVS infection (**Figure 3A—C**). Nonetheless, *Ifng* transcript did not increase significantly in these two T cell subsets over time (**Figure 3D**). NK, NKT1 and effector CD8^+^ T cells did not elicit a type 3 response. Consistent with this finding, no *Il17a* or *Rorc* transcripts were observed (**Figures 3E&F, S3C&E**). Although NKTfh appear to express type 3 immune signature, they do not express *Il17a* or *Rorc* (**Figures 3E&F, S3C&E**). Taken together, the results thus far indicate that NK and NKT1 cells elicit a type 1 immune response typified by IFN-γ production in response to respiratory LVS infection.

### Respiratory LVS infection enhances MAIT cell responses in the lungs of NKT cell-deficient CD1d^-/-^ mice

We previously reported that NKT cell-deficient *Cd1d*^-*/-*^ mice resistant respiratory infection with 8,000 cfu LVS—the same dose at which ≥50% of C57BL/6 mice succumb^24^. Moreover, MAIT cells expand in naïve *Cd1d*^-*/-*^ mice^69^, and play critical roles during LVS infection^31^. Nonetheless, the mode of MAIT cell activation and the specific MAIT cell subset involved in protection from respiratory LVS infection are unknown. Hence, we investigated MAIT cell involvement in pulmonary immune responses to LVS infection in *Cd1d*^-*/-*^ mice. As reported above, both MAIT1 and MAIT17 cells expanded in response to respiratory LVS infection of C57BL/6 mice (**Figures 1 & 2**). To fully explore the dynamics of MAIT cell involvement in the context of a protective response, single cell transcriptomic analyses were performed on the B220^+^ B cell-negative as well as CD14^+^, CD11b^+^ and CD11c^+^ myeloid cell-negative fraction of CD45^+^ cells flow sorted from the lungs of mock or LVS-infected, *Cd1d*^-*/-*^ mice. Quality control yielded a total of 4,986 high quality and comparable cells across the samples (**Figure 4A**). As in LVS-susceptive C57BL/6 lungs, immune cell accumulation was observed that only began by 5 dpi in TLD-resistant *Cd1d^-/-^*lungs (**Figure 4B&C**). Additional analyses confirmed the presence of a type 1 response driven by NK and CD8 T cells, as well as a type 3 response mainly driven by MAIT and γδ T cells (**Figure 4D&E, S4A-D**). Thus, LVS infection also induces ITL-dominated type 1 and type 3 immune responses in TLD-resistant NKT cell deficient, *Cd1d*^-*/-*^ mice.

**Figure 4:**
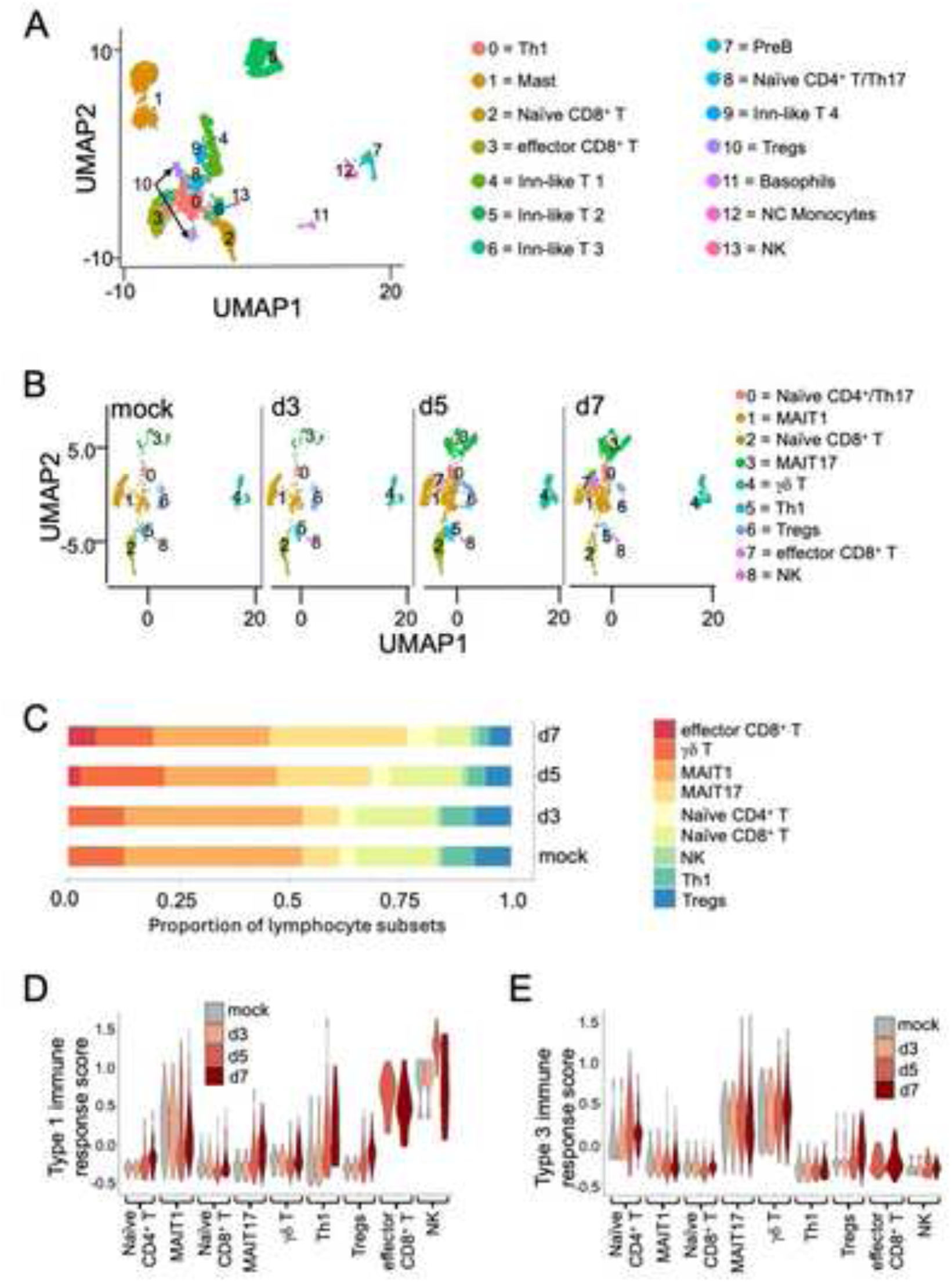
LVS infected NKT cell-deficient CD1d-null mice elicit type 1 and type 3 inflammatory responses from lung innate-like T lymphocytes. Three six-week-old *Cd1d^-/-^* mice were inoculated by the i.n. route with either PBS (mock) or ∼8,500 cfu LVS. At 3, 5, and 7 dpi, myeloid cell and B cell negative fraction of CD45^+^ lymphocytes was flow sorted and subjected to scRNAseq as in figure 2. **A.** UMAP plot of 4,986 myeloid and lymphoid cells obtained post QC by scRNAseq of sorted lung CD45^+^ T cells from mock and LVS infection at 3, 5 or 7 dpi. **B.** UMAP plot of lymphocytes colored by inferred cluster identity and separated by dpi. Over three thousand (3,360) lymphocytes from **A** were in silico extracted and re-clustered for further analyses. **C.** Proportion of lymphocyte subsets by cluster per condition (mock, 3, 5, and 7 dpi). **D.** Violin plot for scaled expression of type 1 immune response gene signature based on genes listed in figure 1E legend. **E.** Violin plot for scaled expression of type 3 immune response gene signature based on genes listed in figure 1F legend. Data for A—E were generated from pooled samples of 3 mice per condition.

To explore the protective features of MAIT cell responses to LVS infection, all MAIT cells from C57BL/6 and *Cd1d^-/-^* cohorts were independently extracted by in silico sorting and subclustered for in-depth analyses. In line with previous reports^69^, at baseline NKT cell-deficient *Cd1d^-/-^* mice exhibited increased total pulmonary MAIT cells compared to C57BL/6 mice (**Figure 5A&B**). Surprisingly, upon LVS infection, C57BL/6 mice showed only a modest accumulation of MAIT cells by 5 dpi, with the most notable accumulation occurring at 7 dpi (**Figure 5A, S5A**). By contrast, LVS-infected *Cd1d^-/-^* mouse lungs exhibited enhanced MAIT cell accumulation by 5 dpi (**Figure 5B, S5B**). Interestingly, we noted that cluster 2—referred to as transitional MAIT cells, which maintained a highly active type 1 inflammatory signature, was distinctly replaced by cluster 1—referred to as hybrid MAIT cells, which largely maintained a type 3 inflammatory signature. Furthermore, hybrid MAIT cells significantly up-regulated the *expression* of *Ikzf3* encoding Aiolos as well as its target genes *Irf8 and Myc* (**Figure S5C**), which are markers of T cell plasticity^70, 71^. C57BL/6 MAIT cells displayed significant levels of both type 1 and type 3 gene expression signatures by 7 dpi (**Figure 5 C&D, S5D&E**). On the other hand, *Cd1d^-/-^* lungs exhibited about a two-fold increase in expression of both type 1 and type 3 gene signatures by 5 dpi, which persisted through 7 dpi (**Figure 5E&F, S4E&F**). The transitional MAIT cell cluster were so called because they decreased over time commensurate with increased MAIT17 cells, which also showed a hybrid MAIT cell gene signature. We hypothesize that transitional MAIT cells may be transitioning into hybrid MAIT and MAIT17 cells. We also observed that, in addition to a predominant type 3 profile, the hybrid MAIT cell cluster maintained some type 1 gene expression signature (**Figure 5B,D&F**). In accordance with previous findings^43^, MAIT cell clusters from both C57BL/6 and *Cd1d*^-/-^ lungs could be distinctly grouped into *Klrg1* and *Il7r* (encoding CD127—the IL-7 receptor)-expressing clusters (**Figure S5F&G**). *Klrg1* was largely expressed by MAIT1 cells, whereas *Il7r* was mostly expressed by MAIT17 cells (**Figure S5F&G**). Taken together, LVS infection distinctly induces enhanced and accelerated type 1 and type 3 MAIT cell responses in TLD-resistant, *Cd1d*^-*/-*^ mice.

**Figure 5:**
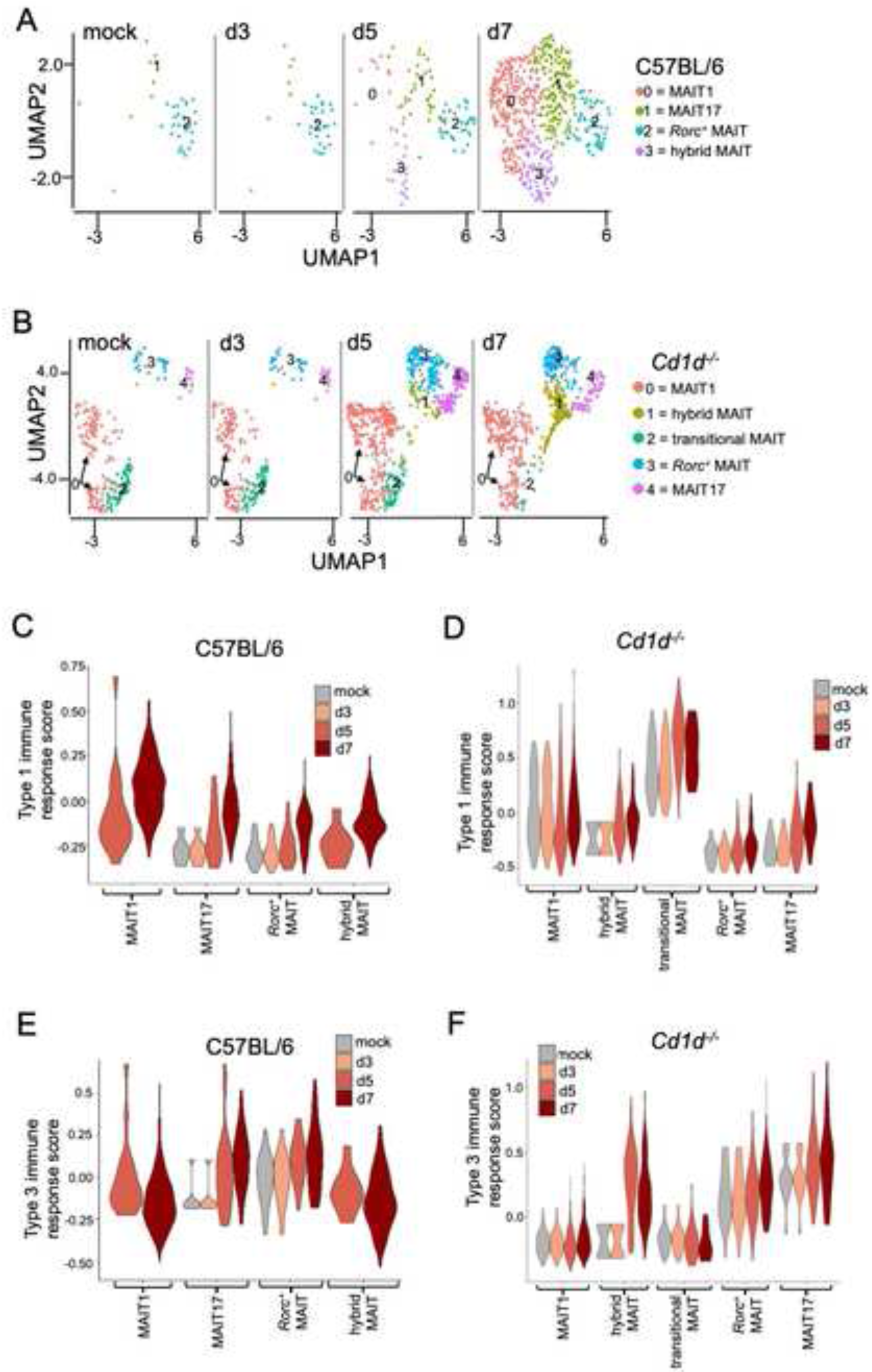
LVS infection elicits an enhanced MAIT cell response in *Cd1d^-/-^* mice. **A.** UMAP plot of pulmonary MAIT cells from C57BL/6 mice colored by cluster identity and separated by condition. Over seven hundred (741) MAIT1 and MAIT17 cells were in silico extracted from figure 2B, re-clustered and annotated for comparative analysis. **B.** UMAP plot of in silico extracted MAIT cells from *Cd1d^-/-^* mice colored by cluster identity and separated by condition. Over sixteen-hundred (1,696) MAIT1 and MAIT17 cells in figure 4B were extracted in silico, re-clustered and annotated for comparative analysis. **C—D.** Violin plot for scaled expression of type 1 inflammatory immune response gene signature from C57BL/6 (**C**) or *Cd1d^-/-^* (**D**) MAIT cells based on genes listed in figure 1E legend. **E—F.** Violin plot for scaled expression of type 3 inflammatory immune response gene signature from C57BL/6 (**E**) or *Cd1d^-/-^*(**F**) MAIT cells based on genes listed in figure 1F legend. Data for A—F were generated from pooled samples of 3 mice per condition.

### LVS infection expands protective lung-resident MAIT cells via TCR & cytokine-mediated mechanisms

Akin to NKT cells^29, 72^, MAIT cells can be activated via TCR-dependent but cytokine-independent, TCR-dependent plus cytokine-dependent, and TCR-independent but cytokine-dependent mechanisms^33, 73–76^. Nur77—encoded by *Nr4a1*, is induced upon TCR stimulation in conventional T, NKT and MAIT cells and, hence, serves as a proxy for T cell activation upon antigen/ligand/agonist recognition^65, 77–82^. Mouse MAIT cell activation by LVS in vitro is MR1-dependent and, hence, by extension, TCR-dependent^31^. By contrast, despite the presence of a functional riboflavin synthesis pathway that is essential for the generation of MAIT cell agonist 5-OP-RU (5-(2-oxopropylideneamino)-6-d-ribitylaminouracil), *M. bovis* infection drives a predominant type 3 MAIT cell response in a MR1-independent manner^83^. Hence, we evaluated the induction of *Nr4a1* and *Il18r* transcripts in pulmonary MAIT cells in vivo during LVS infection of both C57BL/6 and *Cd1d*^-*/-*^ mice. All MAIT cell subsets in C57BL/6 mice significantly upregulated *Nr4a1* expression by 7 dpi. All MAIT cells also rapidly (as early as 5 dpi) upregulated *Il18r1* expression (**Figure 6A**). Similarly, MAIT cell clusters in LVS infected *Cd1d*^-*/-*^ lungs significantly upregulated both *Nr4a1* and *Il18r* expression, but both transcripts were upregulated by 5 dpi and sustained through 7 dpi. Interestingly, the transitional MAIT cell cluster, which was not detected in LVS-infected C57BL/6 mice, displayed reduced *Nr4a1* expression and contained cells in which *Il18r1* was upregulated by 7 dpi (**Figure 6B**).

**Figure 6:**
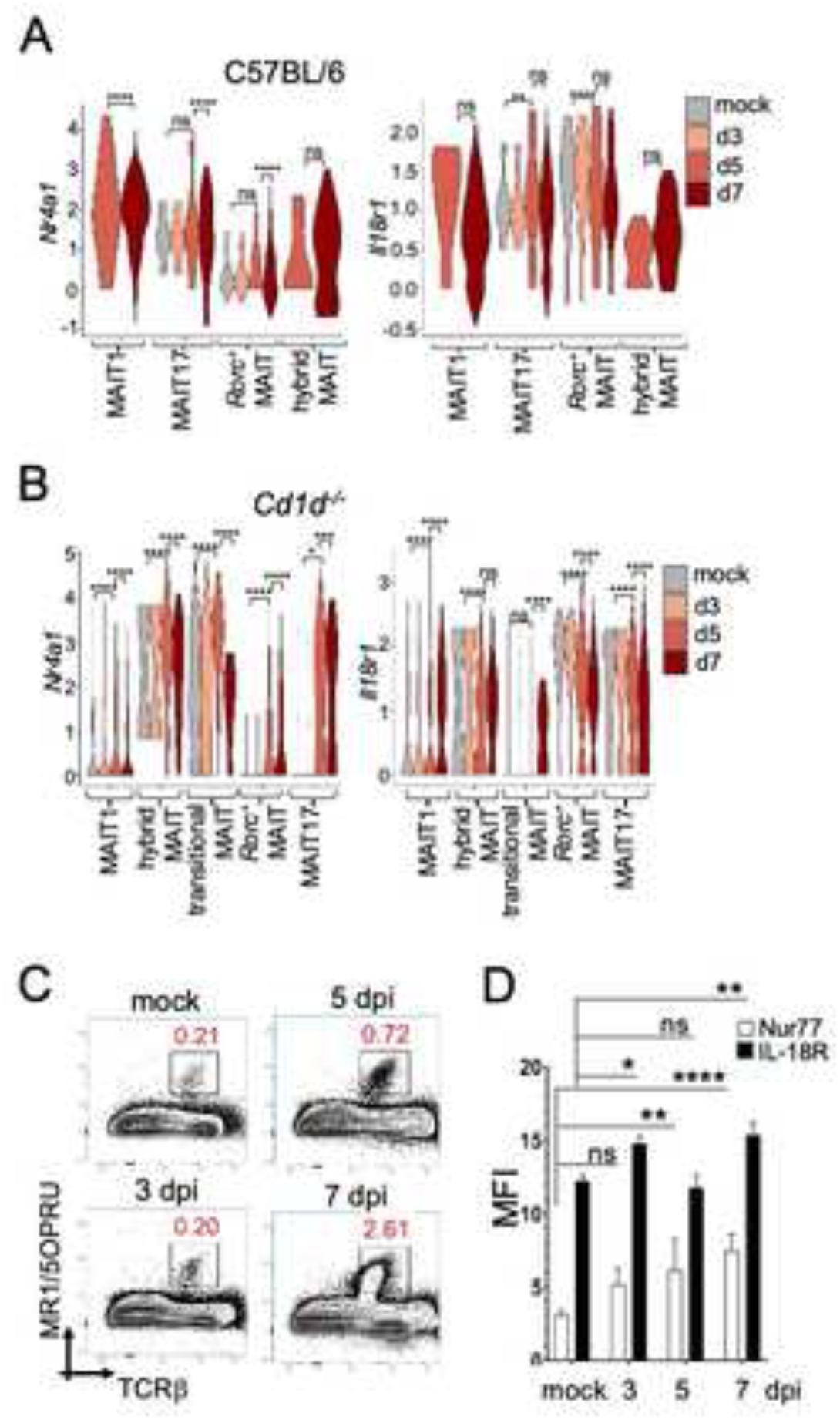
LVS infection expanded lung-resident MAIT cells via TCR & cytokine-mediated mechanisms. **A.** Violin plots for *Nr4a1* and *Il18r1* expression in C57BL/6 pulmonary MAIT cell clusters. **B.** Violin plots for expression of *Nr4a1* and *Il18r1* respectively in *Cd1d^-/-^*pulmonary MAIT cell clusters. **A&B** were generated from pooled samples of 3 mice per condition. Mann Whitney test with Bonferroni posttest correction. *p<0.05, **p<0.01, ***p<0.001, ****p<0.0001. **C—D.** Representative flow plots (**C**) and MFI normalized to isotype control (**D**) quantifying Nur77 and IL18Ra protein levels in pulmonary MAIT cells in mock or LVS infected mice at 3, 5, and 7 dpi. Bars represent the mean+SEM (*n*=3—5 mice per group per experiment). Two-way ANOVA with Tukey’s multiple comparison test. *p<0.05, **p<0.01, ***p<0.001, ****p<0.0001.

Next, the modes of MAIT cell activation were evaluated by flow cytometry in LVS-infected Nur77-reporter *Nr4a1^eGFP^* mice. *Nr4a1^eGFP^* mice significantly accumulated MAIT cell by 5 dpi with continued accumulation at 7 dpi (**Figure 6C**). eGFP expression, and by proxy Nur77 protein levels, significantly increased in pulmonary *Nr4a1^eGFP^* MAIT cells as early as 5 dpi and were sustained through 7 dpi (**Figure 6D**). Concurrently, there was significant increase in IL18Ra protein in *Nr4a1^eGFP^* MAIT cells (**Figure 6D**), indicating a combined TCR and cytokine mediated activation of MAIT cells. Hence, both sustained TCR and cytokine mediated activation elicited MAIT cell function during LVS infection in NKT cell sufficient and deficient mice.

### Enhanced MAIT17 cell response protects LVS-susceptible, immunodeficient *RAG2^-/-^* mice from severe morbidity & mortality caused by respiratory LVS infection

We next sought to compare the expression of hallmark MAIT1 and MAIT17 cytokine genes in lungs of mock and LVS infected C57BL/6 and *Cd1d^-/-^* mice. In the C57BL/6 lungs, distinct MAIT1 and hybrid MAIT cell clusters expressed significant levels of *Ifng* and *Tbx21* by 7 dpi (**Figure 7A**). Curiously, unlike NK and NKT1 cells in mock infected C57BL/6 lungs (**Figure 3D**), naïve MAIT1 cells in the lungs of infected C57BL/6 mice have not transcribed their *Ifng* and *Tbx21* loci (**Figure 7A**). By contrast, in the *Cd1d*^-/-^ lungs, naïve MAIT1, transitional MAIT and hybrid MAIT cell clusters have already transcribed both the *Ifng* and *Tbx21* loci. Further, LVS infection of *Cd1d*^-/-^ lungs augmented the levels of *Ifng* and *Tbx21* transcripts by 5 dpi that were sustained until 7 dpi (**Figure 7B**). Surprisingly, however, a small fraction of MAIT17 cells in the C57BL/6 lungs had transcribed the *Ifng* and *Tbx21* loci, whereas those in *Cd1d*^-*/-*^ lungs did not (**Figure 7A&B**). Interestingly, a mixed Th1/Th17-like MAIT cell cytokine profile was previously reported after in vitro stimulation of MAIT cells^42^. Moreover, in the C57BL/6 lungs, MAIT17 and hybrid MAIT cell clusters expressed *Il17a* and *Rorc*, which codes for RORγt—a Th17-typic transcription factor that drives type 3 responses, only by 5 or 7 dpi with LVS (**Figure 7C**). Similar to C57BL/6 lungs, naïve MAIT cells have not transcribed *Il17a,* although naive *Rorc*^+^ MAIT and MAIT17 clusters have already transcribed the *Rorc* locus. By 5 dpi, *Il17a* and *Rorc* transcripts were significantly increased and persisted through 7 dpi within the MAIT17 and hybrid MAIT cell clusters (**Figure 7D**). Again, a small fraction of MAIT1 cells in the C57BL/6 lungs had transcribed the *Il17a* and *Rorc* loci, which were not found in the *Cd1d*^-*/-*^ lungs (**Figure 7C&D**). Last, the expression of other hallmark type 1 and type 3 cytokine genes followed similar patterns (**Figure S6A&B**).

**Figure 7:**
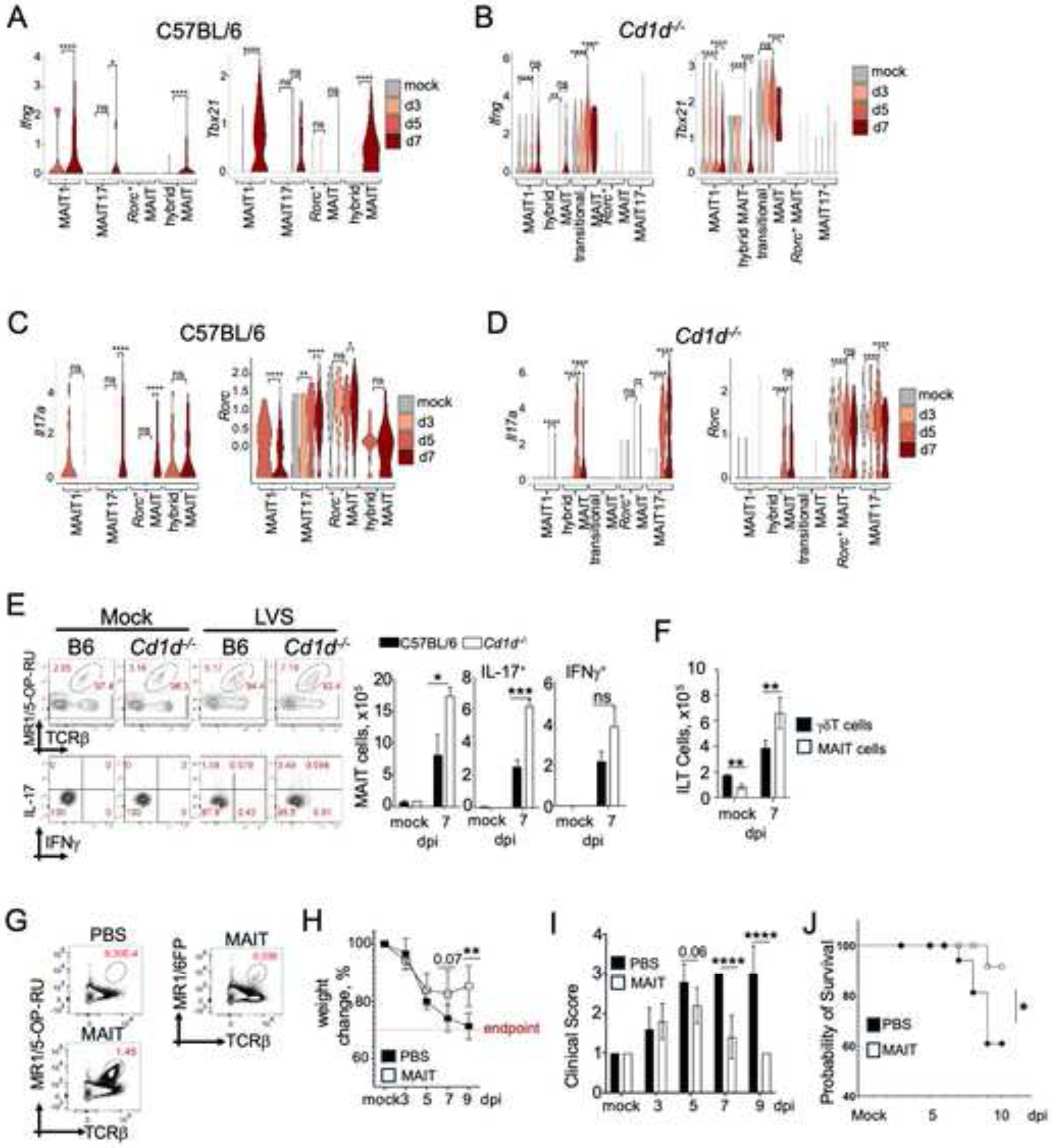
MAIT17 cells protect mice from severe disease post pulmonary LVS infection. **A.** Violin plots for expression of *Ifng* and *Tbx21* in C57BL/6 pulmonary MAIT cell clusters. **B.** Violin plots for expression of *Ifng* and *Tbx21* in *Cd1d^-/-^* pulmonary MAIT cell clusters. **C.** Violin plots for expression of *Il17a* and *Rorc* in C57BL/6 pulmonary MAIT cell clusters. **D.** Violin plots for expression of *Il17a* and *Rorc* in *Cd1d^-/-^* pulmonary MAIT cell clusters. Data for A—D were generated from pooled samples of 3 mice per condition. Mann Whitney test with Bonferroni posttest correction. *p<0.05, **p<0.01, ***p<0.001, ****p<0.0001. **E.** Representative flow plots and quantification of pulmonary MAIT cells in C57BL/6 and *Cd1d^-/-^* mice. Total MAIT cells were identified and enumerated by co-staining with MR1/5-OP-RU tetramer and anti-mouse TCRβ antibody, followed by intracellular staining for IL17A and IFN-γ in lung MAIT cells of mock or LVS-infected C57BL/6 and *Cd1d^-/-^*mice at 7 dpi. MR1-6FP tetramer was used as a control. Bars represent the mean+SD (*n*=5 mice/group per experiment from 3 independent experiments). Two-way ANOVA with Bonferroni posttest correction. *p<0.05, **p<0.01, ***p<0.001. **F.** Quantification of pulmonary MAIT and γδ T cells in C57BL/6 and *Cd1d^-/-^* mice. Bars represent the mean+SEM (*n*=5 mice/group per experiment; representative of at least 3 independent experiments). Two-way ANOVA with Tukey’s multiple comparison test. *p<0.05, **p<0.01, ***p<0.001. **G.** Representative flow plots of pulmonary MAIT cells in *RAG2^-/-^* mice post transfer and mock or LVS infection. MAIT cells were flow sorted from LVS-inoculated *Cd1d^-/-^* mice at 7 dpi and 80,000-100,000 (MR1-5OPRU^+^) cells transferred into each lymphopenic *RAG2^-/-^*mice (*n*=5 mice/experiment), After ∼18 hours, mice were inoculated by the i.n. route with ∼8,000 cfu LVS. Controls received PBS (*n*=5 mice/experiment). At 10 dpi, MAIT cells were tracked in all mice. **H&I.** Morbidity and mortality of LVS-infected *RAG2^-/-^* mice post PBS administration or MAIT cell transfer. Weight loss (**H**) and clinical disease scores (**I**) were monitored daily but only those for 3, 5, 7 and 9 dpi are shown. Bars represent the mean+SEM; representative plots of 3 independent experiments. *n*=5 mice per group per experiment. Student unpaired t test *p<0.05, **p<0.01, ***p<0.001, ****p<0.0001. **J.** Survival of *RAG2^-/-^* mice after infection with 8,000 cfu LVS post PBS administration or MAIT cell transfer. Kaplan-Meier survival analysis; *n*=15 mice per group per experiment. Cumulative data from 3 independent experiments; Log-Rank (Mantel-Cox) test; *p<0.05.

The presence of *Ifng* and *Il17a* transcripts, albeit at low levels, in naïve MAIT cells of *Cd1d*^-*/-*^ lungs suggested that these cells are poised to respond quickly to stimulatory signals from LVS infection, which is distinct from naïve MAIT cells of C57BL/6 lungs in which *Ifng* and *Il17a* transcripts were absent. Hence, we ascertained the dynamics of cytokine protein production by MAIT cells in response to LVS infection by intracellular cytokine staining and flow cytometry (**Figure 7E**). As previously established^69^, we found higher frequency of pulmonary MAIT cells (TCRβ^+^ MR1/5-OP-RU^+^) at baseline in *Cd1d*^-*/-*^ mice and concordantly increased cell numbers by 7 dpi when compared to C57BL/6 mice (**Figure 7E**). Accordingly, a subset of RORγt^+^ MAIT cells increased in numbers over time which peaked at 7 dpi with LVS (**Figure S6C**). Consistent with the increased MAIT cells was the significantly increased IL-17A-producing MAIT cell accumulation in *Cd1d*^-*/-*^ mouse lungs when compared to C57BL/6 mouse lungs at 7 dpi (**Figure 7E**). A similar trend was observed with IFN-γ-producing MAIT cells, but the difference was not statistically significant (**Figure 7E**). These results indicate that, in addition to a comparable MAIT1 response, TLD-resistant lungs exhibit increased accumulation of IL-17A-producing MAIT cells upon LVS infection.

γδ T cells also surveil the lung mucosa and play roles in infection and immunity^84^. Further, γδ T cells have been suggested as a source of IL-17A that protects against respiratory LVS infections^48^. Hence, comparative analyses of γδ T and MAIT cells were conducted. As previously reported^85^, there were more γδ T cells in *Cd1d*^-*/-*^ mice as compared to C57BL/6 mice. Accordingly, these cells expanded to significantly high numbers and produced both IFN-γ and IL17A by 7 dpi in *Cd1d*^-*/-*^ lungs when compared with C57BL/6 lungs (**Figure S6D**). Moreover, NKT cell deficient, *Cd1d*^-*/-*^ mice have increased levels of pulmonary γδ T cells than MAIT cells at baseline. LVS infection induced significant expansion of total γδ T and MAIT cells in *Cd1d*^-*/-*^ mouse lungs 7 dpi with LVS. Nonetheless, significantly more MAIT cells than γδ T cells accumulated by 7 dpi (**Figure 7F**). Despite disproportionate γδ T and MAIT cell expansion, there was no significant difference in either IL-17A-or IFN-γ-producing γδ T and MAIT cells at 7 dpi (**Figure S6E**). Thus, MAIT cells are significant contributors to IL-17 production in the lungs post LVS infection.

Previous work showed that the absence of MAIT cells renders mice susceptible to severe morbidity due to pulmonary LVS infection. Thus, MAIT cells are essential for protective responses to LVS infection^31, 44, 86^. Whether these cells are sufficient to protect from LVS infection-induced morbidity remained as yet unknown. To that end, 80,000—100,000 MAIT cells were flow sorted from LVS-infected *Cd1d*^-*/-*^ mice at 7 dpi and adoptively transferred intravenously into immune deficient *RAG2^-/-^* mice. LVS-infected *Cd1d*^-*/-*^ mice were used as donors because MAIT17 cells predominate the lungs of infected *Cd1d*^-*/-*^ mice (**Figure 7E**). Control *RAG2^-/-^* mice received PBS. Eighteen hours post MAIT cell transfer or PBS treatment, recipient and treated *RAG2^-/-^* mice were challenged with ∼8,000 cfu (LD_50_) LVS by the intranasal route. We found that by 9 dpi, the transferred MAIT cells were detectable in the lungs of *RAG2*^-/-^ mice that received cells but not PBS (**Figure 7G**). Consistent with this finding, MAIT cell-recipient *RAG2*^-/-^ mice showed reduced weight loss (**Figure 7H**), lower disease manifestation (**Figure 7I**) and improved survival (**Figure 7J**) compared to control mice that received PBS. Taken together, the data suggest that MAIT17 cells, which predominate LVS infected *Cd1d*^-*/-*^ mice, are sufficient to confer protection from morbidity and mortality caused by LVS infection of mice.

## DISCUSSION

The goal of this study was to understand the dynamics of innate-like lymphocyte responses during the onset of tularemia-like disease (TLD) in a mouse model of LVS infection. For this, a transcriptomic map of lymphocytes isolated from the lungs of TLD-sensitive C57BL/6 and TLD-resistant CD1d-null mice was developed at single cell resolution. A comparative study of the two transcriptomic maps and confirmatory immunoassays revealed that both strains elicited a type 1 and type 3 inflammatory response. NK, NKT1 and MAIT1 cells were responsible for the type 1 inflammatory response and MAIT17 and γδ T cells for the type 3 response. The TLD-resistant mouse strain distinguished itself with a higher number of MAIT cells over the TLD-sensitive mouse strain, even during homeostatic conditions, as reported previously^66^. Out of this skewed MAIT cell enriched *Cd1d*^-*/-*^ mouse emerged an IL-17-dominated response, which upon transfer into immunodeficient *RAG2^-/-^* mice conferred protection against the development of TLD. Collectively, these findings position MAIT cells as potential mediators of interleukin-17-dependent protection from pulmonary TLD. These findings significantly move the field forward in the following ways: First, they support a sequential type 1 to type 3 inflammatory response model to confer protection in the acute phase of TLD. Second, as further discussed below, the data implicate an iBALT-mediated mechanism of protection. And last, they inform a MAIT17-targeted vaccine design.

A past study implicated IFN-γ-producing MAIT1 cells as the key mediators of protection from systemic LVS infection^44^. That report did not find MAIT17 cells, which is not surprising as MAIT cells are rare, if at all present, in the spleen and the circulation and, hence, appears at odds with other studies which have revealed a critical role for IL-17 in resistance against pulmonary TLD^46–50^ and with the current report. Further, one of the studies pin-pointed γδ T cells as the main source of IL-17 and the key mediator of resistance to TLD^48^. The current study suggests that both IFN-γ-producing NK, NKT1 and MAIT1 cells and IL-17-producing MAIT17 and γδ T cells are sequentially involved in the early—3–5 dpi, and late—5–7 dpi, phases of LVS infection, respectively. This sequential response appears to confer resistance against TLD even before the induction of adaptive immune responses by conventional T cells. This model of resistance to LVS infection-induced TLD explains the vast literature (discussed in the Introduction) in which the absence of early IFN-γ-producing innate-like lymphocytes—NK, NKT1 and MAIT1 cells, leads to TLD even though IL-17-producing MAIT17 and γδ T cells are present. We reason that the early IFN-γ response keeps the intracytosolic LVS burden and dispersal under control, as observed in our previous comparative study between TLD-sensitive C57BL/6 and TLD-resistant *Cd1d^-/-^* mouse responses to LVS infection^24^. Further, the induction of IL-17 in the later phase of the acute infection is essential for LVS clearance. This IL-17 response does not kick in early enough to control the infection when the host is relieved of the prostaglandin E2-induced immune suppression by the bacterium and, hence, requires the early IFN-γ response. But the persistence of an oversized runaway type 1 inflammatory response is detrimental as we reported previously^24^. In support of this notion is the finding herein where MAIT cells are yet to open their cytokine gene loci in the TLD-sensitive C57BL/6 mice but have already done so at steady state in the TLD-resistant *Cd1d^-/-^* mice. While several innate and adaptive immune mechanisms are perhaps involved in IL-17-mediated LVS clearance, how this is accomplished requires further investigation.

One potential IL-17-mediated mechanism may involve the induction of the tertiary lymphoid structures (TLSs) called the iBALT in the lungs of LVS-infected mice. Our previous study revealed that TLD-sensitive C57BL/6 mice poorly develop iBALTs in response to respiratory LVS infection. By contrast, the induction of iBALT was a key distinguishing feature of the TLD-resistant *Cd1d^-/-^* mice. Thus, when combined with another study^87^, NKT cells appear to inhibit iBALT formation. The role of MAIT cells in iBALT formation, however, remains unclear. MAIT cells, like other ITLs, are present in TLSs where they are thought to enhance a return to homeostasis post insult^88, 89^. Moreover, IL-17 is required for TLS formation^90, 91^. As seen in this report, MAIT17 cells produce a significant amount of IL-17. Consistent with these findings, a preliminary analysis of our current scRNAseq data showed the expression of several genes (*Cxcr4, Bcl6, Lta, Tnfsf14, Il17a, Tnf,* and *Il22*) implicated in TLS and/or iBALT formation^92, 93^. We found a modest upregulation of TLS-promoting factors in C57BL/6 MAIT cells by 7 dpi. By contrast, there was remarkably higher expression of iBALT-promoting factors in *Il17a*-expressing MAIT cells in LVS-infected *Cd1d^-/-^* lungs by 5 dpi that persisted through 7 dpi. Furthermore, a higher frequency of *Il17a*-expressing MAIT cells from *Cd1d^-/-^* lungs expressed elevated levels of iBALT-promoting factors compared to C57BL/6 MAIT cells (data not shown). These results suggest that MAIT17 cells may contribute to TLS formation to protect mice from morbidity caused by respiratory LVS infection. Additional studies shall help firm this conclusion.

We found that LVS infection first induced *Nr4a1* expression—a proxy for TCR engagement and cell activation^66^, and eventually *Il18r1* expression in NKT, MAIT1 and MAIT17 cells. This finding suggests that vaccine design against *F. tularensis* should consider adjuvants that induce both TCR- and cytokine-dependent MAIT cell activation. Moreover, a recent study indicated that the TCR-dependent pathway of MAIT cell activation is critical, as the *ribD* gene of type A *F. tularensis* contains mutations that render them defective in riboflavin biosynthesis. Consequently, the type A strain lacks the MAIT cell antigen 5-OP-RU. This *ribD* mutation when introgressed into LVS, renders it incapable of TCR-dependent MAIT cell activation as well^94^. Hence, the inability to activate MAIT cells may explain the extreme virulence of type A *F. tularensis*. We found that the lack of NKT cells in *CD1d^-/-^* mice tempered this virulence^24^. Hence, a type A vaccine may consider deletion of the NKT cell TCR agonist and the addition of a functional *ribD* gene. Engineering these changes into SchuS4—a prototype type A *F. tularensis* strain, may tame its extreme virulence to provide an effective homotypic vaccine. Alternatively, the deletion of the gene/s that encode the NKT cell agonist in LVS may make it a more reliable, effective vaccine. Thus, the findings in this report explain how the sequential type 1 to type 3 inflammatory response can lead to protective immunity to one of the most virulent pulmonary pathogens that involves the induction of iBALTs and how these features can be harnessed for vaccine design.

## Acknowledgements

We greatly appreciate the technical assistance provided Ms. W. Hu, Dr. Y. Su and Ms. A.J. Joyce. SJ is a Research Career Scientist of the US Department of Veterans Affairs supported by IK6 BX004595. GDO and NUC were supported by T32GM007347 and NUC by F30HL159941. This work was supported by Merit Awards BX001444 & BX001610 to SJ and BX000915 to HMSA as well as by NIH grants AI137082 to SJ, AI139046 to LVK, and HL136664 to DCN. Flow Cytometry Shared Resource is supported by CA68485 & DK058404. VANTAGE Shared Resource is supported by RR024975, CA68485, EY08126 & RR030956.

## Author contributions

GDO performed the experiments, analyzed the data, and wrote and edited the manuscript. AK co-conceived the project, helped to write the AI137082 proposal, performed experiments, analyzed the data, and edited the manuscript. FG assisted with several experiments. NUC assisted with some flow cytometry experiments. LW assisted with some of the experiments and edited the manuscript. DCN supervised some flow cytometry experiments and flow data analyses. LVK assisted with data analysis, provided critical reagents, and edited the manuscript. HMSA co-conceived the project, assisted in data analysis, and edited the manuscript. SJ conceived the project, sought funding, supervised the project, analyzed the data, and edited and finalized the MS.

## STAR METHODS

### RESOURCE AVAILABILITY

#### Lead Contact

Further information and reasonable requests for resources and reagents should be directed to and will be fulfilled by the lead contact, Sebastian Joyce.

#### Materials Availability

This study did not generate new unique reagents or mouse lines.

#### Data and Code Availability

Single-cell RNA-seq data have been deposited at NCBI Sequence Read Archive (SRA) under the accession numbers SAMN40657083— SAMN40657092 (see Key Resource Table) and are publicly available as of the date of publication. The current study does not report original code(s). All flow cytometry and infection metrics data reported in this paper will be shared by the lead contact upon request. Any additional information required to reanalyze the data reported in this paper is available from the lead contact upon request.

### EXPERIMENTAL MODEL DETAILS

#### Mice

Age-matched, 6-to-12-week-old male mice were used for experiments described herein.

Prior epidemiological and experimental reports on *F. tularensis* have indicated that males were more susceptible to infection than females^95–101^. C57BL/6J (RRID:MGI:3028467), Nur77^GFP^ (C57BL/6-Tg(Nr4a1-EGFP/cre)820Khog/J; RRID:MGI:5007652), and *RAG2^-/-^* (B6.Cg-Rag2tm1.1Cgn/J; RRID:IMSR_JAX:008449) mice were purchased from the Jackson Laboratory (Bar Harbor). *Cd1d^-/-^* (B6(C)-Cd1d1tm1.2Aben/J; RRID:IMSR_JAX:017294) mice have been described^24^. Mice were bred and maintained in the VUMC (Vanderbilt University Medical Center) vivarium and provided with food and water ad libitum. LVS infection was performed in an ABSL-2 facility. Vanderbilt’s IACUC approved the experiments described here. The current study did not generate any unique mouse strains. All mouse strain maintained by us for this study are available to investigators from the lead contact upon request.

#### Study approval and Ethics Statement

All procedures for use of mice in the study were approved by the Institutional Animal Care and Use Committee at Vanderbilt University School of Medicine and VUMC. Anesthesia was performed using 1–5% isoflurane or 100 mg/kg and 10 mg/kg ketamine and xylazine, respectively. Euthanasia was performed by CO_2_ overdose followed by cervical dislocation.

#### Bacterial infection

*F*. *tularensis* LVS was provided by S. Khader (Washington University, St. Louis, MO). Preparation of working stocks and cfu determination from infected tissue were performed as described^102^. LD_50_ was determined by the method of Reed and Muench^103^. All mice used in infection experiments were blinded prior to anesthesa and LVS inoculation. Male, 6-to-12-week-old mice were anesthetized by i.p. administration of a ketamine/xylazine mixture and ∼8–10×10^3^ cfu LVS were administered i.n. in 50 μL sterile PBS. Mice were monitored daily for weight loss and signs of morbidity. Criteria for clinical score were developed based upon observation of mice from at least three separate experiments: 1, no outward signs of illness; 2, consistently ruffled fur; 3, hunched back and altered gait; and 4, reduced mobility/reaction to stimulus, labored breathing, lethargy. Mice were humanely euthanized when weight loss exceeded 30% of the weight at the start of the experiment.

### METHOD DETAILS

#### Tissue processing

Spleen and lungs were processed as previously described^104, 105^. Briefly, mice were euthanized with CO_2_ and lungs were perfused with PBS. Lungs were minced and incubated in medium containing Collagenase/DNAse for 45 minutes at 37°C sometimes with vigorous agitation at 15-minute intervals. Lung tissue was pushed through a 70 μm nylon screen to obtain a single-cell suspension. Red blood cells were lysed, and the cells were resuspended in PBS containing 2% FBS and 50nM Dasatinib. Cells were further filtered through 40 μm nylon screen before downstream use. Both left and right lung lobes were used for each indicated experiment. For intravascular vs resident T cell measurements, 2 μM FITC-conjugated anti-mouse CD45.2 antibody was retro-orbitally injected into each mouse 3-5 minutes before culling.

#### Flow cytometry

NKT and MAIT cells were analyzed as previously described using respective NKT- and MAIT-specific tetramers^105^. All data were acquired on Canto II 3-Laser 8-color flow cytometer or 4-laser Cytek Aurora spectral cytometer and analyzed with FlowJo software (FlowJo, LLC). Cell populations were identified as follows: B (B220^+^), T (TCRβ^+^ and/or CD3ε^+^), NKT (TCRβ^+^CD3ε^+^CD1d/αGC tetramer^+^), MAIT (TCRβ^+^CD3ε^+^MR1/5-OP-RU tetramer^+^), and γδ T cells (TCRγδ^+^CD3ε^+^) cells. NKT and MAIT subsets were identified by evaluating intracellular staining for hallmark cytokines (IL-17A, IFNγ, IL-4, IL-5, and IL-13), or intranuclear staining for hallmark transcription factors (Tbet, RORγt). Ligand-loaded and -unloaded CD1d and MR1 tetramers were provided by the NIH Tetramer Core Facility. Cell counts were determined using AccuCheck counting beads (Invitrogen) by using manufacturer’s recommended method.

#### Adoptive Transfers

*Cd1d*^-/-^ mice were infected with ∼8—9×10^3^ cfu LVS. Seven dpi, lungs from 3—5 mice were processed, stained, and subsequently sorted for MAIT cells using 4-laser Cytek Aurora cell sorter. Eighty thousand to 100,000 MAIT cells were then transferred by tail vein injection into recipient RAG2^-/-^ mice with subsequent inoculation with LD_50_ LVS 18 hours after transfer. Animals were monitored and MAIT cells characterized as described above.

#### Single cell RNA sequencing

Lung single-cell suspensions were generated from PBS treated or LVS-inoculated mice at 3, 5, 7 dpi as previously described^105^.

#### Single-cell RNA library generation and sequencing

Isolated total lung single-cell suspensions were depleted of dead cells by flow cytometry and similarly enriched for T lymphocytes and subjected to droplet-based massively parallel single-cell RNA sequencing using Chromium Single Cell 5’ (v1) Reagent Kit per manufacturer’s instructions (10x Genomics). Briefly, cell suspensions were loaded at 1,000 cells/μL with the aim to capture 10,000 cells/lane. Each cell in the 10x Chromium Controller generated GEM droplets was labeled with a specific barcode, and each transcript labeled with a unique molecular identifier (UMI) during reverse transcription. The barcoded cDNA underwent 16 cycles of amplification and SPRI bead purification before library generation. Libraries were prepared from amplified cDNA and target enrichment products by fragmentation, end repair, A-tailing, adapter ligation, and sample index PCR per manufacturer’s instructions. Libraries were sequenced on a NovaSeq S4 (200 cycle) flow cell, targeting 50,000 read pairs/cell.

#### Single-cell RNA-seq data processing

The raw gene expression matrices were generated by the Cell Ranger software (10x Genomics, version 7.0.1) available on the 10x website. After demultiplexing, the resulting fastq files were aligned against the mouse reference genome mm10 with cellranger count. The output filtered gene count matrices were analyzed by R software (v.3.5.3) with the Seurat^106^ package (v.3.0.0). Cells that had less than 200 and more than 5000 detected genes were filtered out. The threshold was chosen for mitochondrial genes as follows: had less than 5% of mitochondrial genes for all samples. Samples with similar conditions were merged, data normalized with default parameters, and most variable genes were detected by the FindVariableFeatures function (FDR < 0.05 cutoff). All samples were combined using Seurat functions, FindIntegrationAnchors and IntegrateData. ScaleData was used to regress out number of UMI’s and mitochondrial content and principal component analysis (PCA) was performed with RunPCA. The UMAP dimensionality reduction was performed on the scaled matrix using the first 20 PCA. For clustering, the FindNeighbors (20 PCA) and FindClusters (resolution 0.4) functions were used. FindAllMarkers was used to compare a cluster against all other clusters to identify the marker genes. For each cluster, the minimum required average log fold change in gene expression was set to 0.25 and the minimum percent of cells that must express gene in either cluster was set to 25%. *Cd1d^-/-^*and B6 samples were analyzed separately but concurrently. The computational platform scType was used for initial cluster annotation^53^. scType combines data from multiple models and organs including lungs to identify a variety of immune and non-immune cells. A marker list adapted from scType and used for identifying each annotated cluster is provided in **supplementary Table 1.** Importantly, using scType, as expected, we did not identify any NKT cell clusters in *Cd1d^-/-^* samples. Cluster annotations were further confirmed manually by verifying signature genes provided in **supplementary Table 1**. To re-analyze lymphocyte sub-populations, we pooled the clusters that were identified as lymphoid in origin and re-ran PCA, tSNE, and clustering to get a better resolution for further analysis. The function FindMarkers was used to compare the cells in a selected cluster across different conditions. The analysis was performed based on the Wilcoxon rank sum test with thresholds average logFC ≥0.25 and adj-pvalue ≤ 0.05 using Seurat. Several R packages were used to generate figures and intermediate data preprocessing, such as ggplot2^107^. Inflammatory signature genes were compiled based on previously published gene signatures^57, 58^. Subsequent analyses of Type 1 cell clusters and MAIT cells were performed as above by pooling annotated clusters.

### STATISTICAL ANALYSIS

Statistical analyses of data were performed using GraphPad Prism version 10.2.2 for Windows, GraphPad Software, San Diego California USA (www.graphpad.com). Where indicated in the figure legends, data were aggregated across several (from two to four) replicate experiments. Otherwise, figures are representative of replicate experiments. Clinical score data was analyzed using Generalized Estimating Equation (GEE) model based on Poisson regression. In the model, clinical score was predicted by group type and day of assessment (dummy coded), and their interaction. We report comparisons between groups at each time of assessment, which were obtained by changing the reference category between days of assessment.

### KEY RESOURCES

#### Reagents

All reagents used for the current report are publicly available. Details including RRID identifiers are provided in the key resources table.

## Key resources table

| REAGENT or RESOURCE | SOURCE | IDENTIFIER |
| --- | --- | --- |
| Antibodies |  |  |
| Ghost dye violet 510 viability dye | Tonbo Biosciences | Cat#: 130870<br>RRID:AB_3101904 |
| BV421 Anti mouse CD14 | Biolegend | Cat#: 123329<br>RRID:AB_2721526 |
| BV421 Anti mouse CD11c | Biolegend | Cat#: 117343<br>RRID:AB_2563099 |
| BV421 anti mouse/human CD11b | Biolegend | Cat#: 101251<br>RRID:AB_2562904 |
| APC/Cy7 anti mouse/human CD45R/B220 | Biolegend | Cat#: 103224<br>RRID:AB_313007 |
| FITC anti mouse/human CD45R/B220 | Biolegend | Cat#: 103206<br>RRID:AB_312991 |
| PECy5.5 anti mouse/human CD45R/B220 | Biolegend | Cat#: 103236<br>RRID:AB_893354 |
| BV605 anti human/mouse CD45R/B220 | Biolegend | Cat#: 103244<br>RRID:AB_2563312 |
| PE anti mouse/human CD45R/B220 | BD Biosciences | Cat#: 553090<br>RRID:AB_394620 |
| PB anti mouse/human CD45R/B220 | Biolegend | Cat#: 103227<br>RRID:AB_492876 |
| APC mCD1d PBS57 | NIH | Cat#: 53840<br>RRID:AB_3101905 |
| APC mCD1d Unloaded | NIH | Cat#: 35070<br>RRID:AB_3101906 |
| BV421 mCD1d PBS57 | NIH | Cat#: 53841<br>RRID:AB_3101907 |
| BV421 mCD1d Unloaded | NIH | Cat#: 27927<br>RRID:AB_3101908 |
| APC MR1 5-OP-RU | NIH | Cat#: 39384<br>RRID:AB_3101909 |
| APC MR1 6FP | NIH | Cat#: 53676<br>RRID:AB_3101910 |
| BV421 MR1 6FP | NIH | Cat#: 65011<br>RRID:AB_3101911 |
| BV421 MR1 5-OP-RU | NIH | Cat#: 65013<br>RRID:AB_3101912 |
| PECy7 anti mouse CD3ε | BD Biosciences | Cat#: 552774<br>RRID:AB_394460 |
| APC anti mouse CD3ε | BD Biosciences | Cat#: 553066<br>RRID:AB_398529 |
| PE anti mouse CD3ε | Biolegend | Cat#: 100308<br>RRID:AB_312673 |
| PE anti human/mouse Tbet | eBioscience | Cat#: 12582580<br>RRID:AB_925762 |
| APC anti mouse Tbet | Biolegend | Cat#: 644814<br>RRID:AB_10901173 |
| PECy7 anti mouse Tbet | Biolegend | Cat#: 644823<br>RRID:AB_2561760 |
| FITC anti mouse CD45.2 | Biolegend | Cat#: 109806<br>RRID:AB_313443 |
| APC anti mouse CD45.2 | Tonbo Biosciences | Cat#: 200454<br>RRID:AB_2621576 |
| BV785 anti mouse CD45.2 | Biolegend | Cat#: 109839<br>RRID:AB_2562604 |
| BV421 anti mouse TCRβ | Biolegend | Cat#: 109230<br>RRID:AB_2562562 |
| APC anti mouse TCRβ | Biolegend | Cat#: 109212<br>RRID:AB_313435 |
| PE Cy7 anti mouse TCRβ | Biolegend | Cat#: 109221<br>RRID:AB_893627 |
| Anti mouse CD16/CD32 (Fc Shield) | Tonbo Biosciences | Cat#: 700161<br>RRID:AB_2621487 |
| PE anti mouse RORγt | BD Biosciences | Cat#: 562607<br>RRID:AB_11153137 |
| PerCPy5.5 anti mouse RORγt | BD Biosciences | Cat#: 562683<br>RRID:AB_2737720 |
| PECF594 anti mouse RORγt | BD Biosciences | Cat#: 562664<br>RRID:AB_2643076 |
| PE anti mouse CD218a (IL-18Ra) | Biolegend | Cat#: 157903<br>RRID:AB_2860732 |
| PE anti mouse PLZF | Biolegend | Cat#: 145804<br>RRID:AB_2561973 |
| PE anti mouse IgG1 | BD Biosciences | Cat#: 554680<br>RRID:AB_395506 |
| Purified mouse IgG1 control | Biolegend | Cat#: 400102<br>RRID:AB_2891079 |
| PerCPy5.5 anti mouse IgG2a | BD Biosciences | Cat#: 550927<br>RRID:AB_10052278 |
| PECy7 anti mouse IFNγ | Tonbo Biosciences | Cat#: 607311<br>RRID:AB_2621871 |
| PE anti mouse IFNγ | BD Biosciences | Cat#: 554412<br>RRID:AB_395376 |
| APC anti mouse IFNγ | BD Biosciences | Cat#: 207311<br>RRID:AB_2621616 |
| FITC anti mouse TCR$\gamma\delta$ | Biolegend | Cat#: 118106<br>RRID:AB_313830 |
| PE anti mouse IL-17A | Biolegend | Cat#: 506904<br>RRID:AB_315464 |
| PECy7 anti mouse IL-17A | Biolegend | Cat#: 506922<br>RRID:AB_2125010 |
| APCCy7 anti mouse IL-17A | Biolegend | Cat#: 506940<br>RRID:AB_2565781 |
| Bacterial and virus strains |  |  |
| *Francisella tularensis* LVS | S. Khader (Lin et al<sup>49</sup>) | doi:<br>10.1016/j.immuni.2009.08.025 |
| Chemicals, peptides, and recombinant proteins |  |  |
| Accucheck Counting Beads | Invitrogen | Cat # PCB100 |
| GolgiPlug Protein Transport Inhibitor | BD Biosciences | Cat # 555029 |
| Brefeldin A | Sigma Aldrich | Cat # B7651 |
| Critical commercial assays |  |  |
| Intracellular Staining Kit | Invitrogen | 00-5523-00 |
| Intranuclear Staining Kit | BD Biosciences | 554714 |
| Deposited data |  |  |
| B6_L_D0 | This Paper | SAMN40657083 |
| B6_L_D7 | This Paper | SAMN40657084 |
| B6_D0 | This Paper | SAMN40657085 |
| Cd1d_D0 | This Paper | SAMN40657086 |
| B6_D3 | This Paper | SAMN40657087 |
| Cd1d_D3 | This Paper | SAMN40657088 |
| B6_D5 | This Paper | SAMN40657089 |
| Cd1d_D5 | This Paper | SAMN40657090 |
| B6_D7 | This Paper | SAMN40657091 |
| Cd1d_D7 | This Paper | SAMN40657092 |
| Experimental models: Organisms/strains |  |  |
| C57BL/6J | Jackson Labs | MGI:3028467 |
| C57BL/6-Tg(Nr4a1-EGFP/cre)820Khog/J | Jackson Labs | MGI:5007652 |
| B6.Cg-Rag2tm1.1Cgn/J | Jackson Labs | IMSR_JAX:008449 |
| B6(C)-Cd1d1tm1.2Aben/J | L. Van Kaer | IMSR_JAX:017294 |
| Software and algorithms |  |  |
| R Studio 2023 12.1+402 | Posit PBC | RRID:<br>SCR_014695 |
| Seurat | Satija Labs<sup>111</sup> | RRID:<br>SCR_007322 |
| Seurat Object | Satija Labs<sup>112</sup> | RRID:<br>SCR_022555 |
| Limma | Ritchie et al<sup>113</sup> | RRID:<br>SCR_010943 |
| ggrepel | Ito & Murphy<sup>114</sup> | RRID:<br>SCR_017393 |
| Dplyr | Varsos et al<sup>115</sup> | RRID:<br>SCR_016708 |
| patchwork | Pedersen<sup>116</sup> | RRID:<br>SCR_000072 |
| tidyverse | Wickham et al<sup>117</sup> | RRID:<br>SCR_019186 |
| Matrix | R-Forge | RRID:<br>SCR_001727 |
| Annotation(Dbi) | Pages et al<sup>118</sup> | RRID:<br>SCR_023487 |
| org.Mm.eg.db | Carlson<sup>119</sup> | RRID:<br>SCR_023488 |
| viridisLite | Garnier et al<sup>120</sup> | RRID:<br>SCR_016696 |
| scatterHatch | Deshpande<sup>121</sup> | https://bioconductor.org/packages/release/bioc/html/scatterHatch.html |
| gridExtra | Auguie & Antonov | RRID:<br>SCR_025249 |
| Presto | Vander Heiden et al<sup>122</sup> | RRID:<br>SCR_013285 |
| RColorBrewer | Cynthia Brewer | RRID:<br>SCR_016697 |
| devtools | Wickham et al<sup>123</sup> | RRID:<br>SCR_016961 |
| EnhancedVolcano | Blighe et al<sup>124</sup> | RRID:<br>SCR_018931 |
| cowplot | Wikel lab | RRID:<br>SCR_018081 |
| msigdb | Liberzon et al,<br>Subramania et al<sup>125, 126</sup> | RRID:<br>SCR_022870 |
| Tempora | Tran & Bader<sup>127</sup> | https://github.com/BaderLab/Tempora |
| Fgsea | Korotkevich et al<sup>128</sup> | RRID:<br>SCR_020938 |
| ReactomeGSA | Griss et al<sup>129</sup> | RRID:<br>SCR_014695 |
| Metap | Dewey<sup>130</sup> | RRID:<br>SCR_014686 |
| scType | lanevski et al<sup>54</sup> | https://sctype.app/<br>https://github.com/lanevskiAleksandr/sc-type |
| Graphpad Prisms 10.2.2 | Graphpad Software | RRID: SCR_002798 |
| FlowJo (version 10.10.0) | FlowJo LLC | RRID: SCR_008520 |

