## Supplementary figures and images for "Single-cell resolution map of innate-like lymphocyte response to *Francisella tularensis* infection reveals MAIT cells’ role in protection from tularemia-like disease"

### Supplementary Figures

Click here to access/download;Supplemental Text and Figures;Figure S6-new.jpg

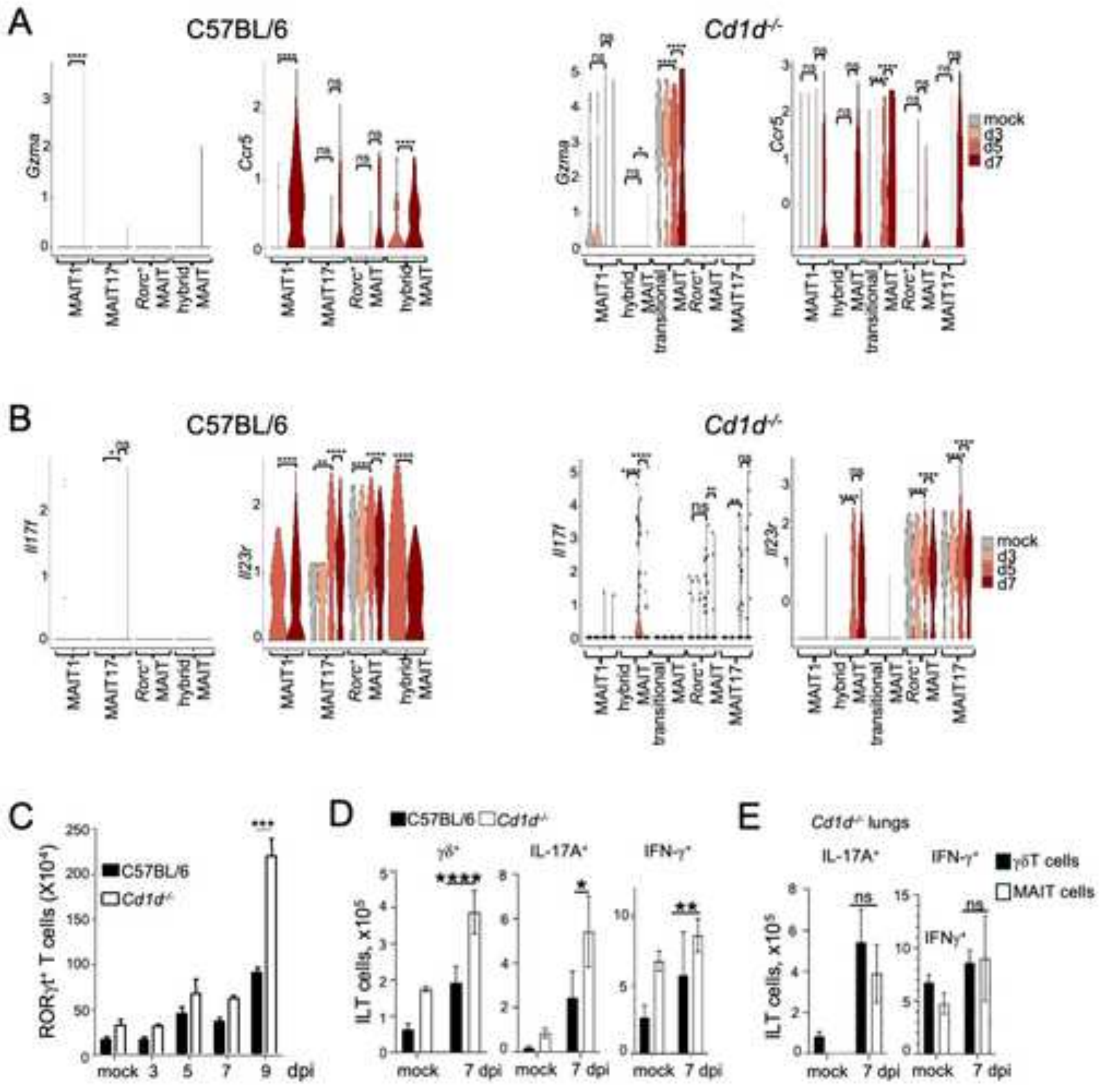


#### Figure S6
